# Slippery flowers as a mechanism of defence against nectar-thieving ants

**DOI:** 10.1101/2020.06.10.144147

**Authors:** Kazuya Takeda, Tomoki Kadokawa, Atsushi Kawakita

## Abstract

**Background and Aims:** The great diversity of floral characters among animal-pollinated plants is commonly understood as the result of coevolutionary interactions between plants and pollinators. Floral antagonists, such as nectar thieves, also have the potential to exert selection on floral characters, but adaptation against floral antagonists has attracted comparatively little attention. We found that the corollas of hornet-pollinated *Codonopsis lanceolata* (Campanulaceae) and the tepals of bee-pollinated *Fritillaria koidzumiana* (Liliaceae) are slippery to nectar-thieving ants living in the plant’s habitat; because the flowers of both species have exposed nectaries, slippery perianths may function as a defence against nectar-thieving ants.

**Methods:** We conducted a behavioural experiment and observed perianth surface microstructure by scanning electron microscopy to investigate the mechanism of slipperiness. Field experiments were conducted to test whether slippery perianths prevent floral entry by ants, and whether ant presence inside flowers affects pollination.

**Key Results:** Scanning electron microscopy observations indicated that the slippery surfaces were coated with epicuticular wax crystals. The perianths lost their slipperiness when wiped with hexane. Artificial bridging of the slippery surfaces using non-slippery materials allowed ants to enter flowers more frequently. Experimental introduction of live ants to the *Codonopsis* flowers evicted hornet pollinators and shortened the duration of pollinator visits. However, no differences were found in the fruit or seed sets of flowers with and without ants.

**Conclusions:** Slippery perianths, most likely based on epicuticular wax crystals, prevent floral entry by ants that negatively affect pollinator behaviour. Experimental evidence of floral defence based on slippery surfaces is rare, but such a mode of defence may be widespread amongst flowering plants.

## INTRODUCTION

The remarkable diversity of floral characters among animal-pollinated plants is commonly understood to be the result of coevolutionary interactions between plants and pollinators. This is exemplified by numerous studies of how the colour, odour and shape of flowers mediate effective attraction of pollinators and their physical contact with anthers and stigmas (e.g. Fenster *et al.* 2004; Papadopulos *et al.* 2013; Johnson and Wester 2017; De Jager and Peakall 2019; Kemp *et al.* 2019). However, flowers are visited by both legitimate pollinators and floral antagonists, the latter including florivores, seed predators, predators of pollinators, and nectar thieves. Such floral antagonists may also exert selective force on floral evolution (Irwin *et al.* 2004; McCall and Irwin 2006; Willmer 2011), but adaptation to such floral antagonists has attracted comparatively little research interest.

Ants generally belong to the floral antagonists (Willmer *et al.* 2009).They are globally ubiquitous and have a major impact on terrestrial ecosystems (Hölldobler and Wilson 1990). A considerable number of plants have independently established mutualisms with ants by offering them rewards and utilising their ecological impact for protection and dispersal; plants in more than 100 families are engaged in defensive mutualisms with ants by producing extrafloral nectar (Weber *et al.* 2015), and plants in more than 80 families exhibit seed dispersal mutualisms with ants (Giladi 2006). However, examples of pollination mutualism involving ants are rare (Dutton and Frederickson 2012). This is thought to be because of their inability to fly, small and hairless body surface (although several ant species are covered with setae (Beattie *et al.* 1984)), frequent grooming behaviour, and chemical compounds on their body that inhibit pollen viability (Beattie *et al.* 1984; Hull and Beattie 1988; Dutton and Frederickson 2012). Nevertheless, ants often visit flowers and consume floral nectar, and are thus mainly regarded as nectar thieves. In addition to consuming nectar, they sometimes attack and deter legitimate pollinators, thereby decreasing pollination success (Galen and Cuba 2001; Tsuji *et al.* 2004; Ness 2006; Lach 2008; Hansen and Müller 2009; Cembrowski *et al.* 2014). Such negative effects of ants are likely to drive the evolution of ant-deterring mechanisms. For example, the floral odours of a wide range of tropical flowers are repellent to ants (Junker and Blüthgen 2008), and several compounds commonly found in floral odour, such as linalool and 2-phenyl ethanol, exhibit such ant repellence (Junker *et al.* 2011). Besides chemical deterrence, extrafloral nectar has been suggested to function as a decoy (Villamil *et al.* 2019), and various floral features have been proposed to act as physical barriers to ants (Willmer *et al.* 2009), but experimental studies to test such functions are limited (e.g. Galen and Cuba 2001; Tagawa 2018; Villamil *et al.* 2019).

*Codonopsis lanceolata* (Campanulaceae) is a perennial vine that produces pendent and campanulate flowers that are almost exclusively pollinated by hornets (Fig. 1A–C). There are five short nipple-shaped spurs on the corners of the corolla base, and the nectar in each spur is often visible as droplets (Fig. 1B). The exposed nature of the nectar suggests that the flower is susceptible to nectar thieving by ants. During preliminary observation, we found that ants visited the flowers and often slipped off the corolla while attempting to walk on it. Further inspections indicated that the abaxial surface and the distal half of the adaxial surface of the corolla of *C. lanceolata* are slippery to ants, whereas the basal half of the adaxial surface, where the pollinators cling to the flower with their forelegs during visitation (Figure 1C), is not slippery (see Figure 2; Supplemental Movie S1, S2). We also discovered slippery perianths in *Fritillaria koidzumiana* (Liliaceae), which similarly bears campanulate flowers and has exposed nectaries. In the present study, we hypothesise that the slippery perianths in *C. lanceolata* and *F. koidzumiana* are a physical defence against nectar-thieving ants and test whether 1) their perianths possess special surface structures that make them slippery to ants (Federle *et al.* 1997; Gaume *et al.* 2002), 2) the slipperiness prevents ants from entering the flowers under natural conditions, and 3) the presence of nectar-thieving ants has a negative effect on pollination.

**Figure 1.**
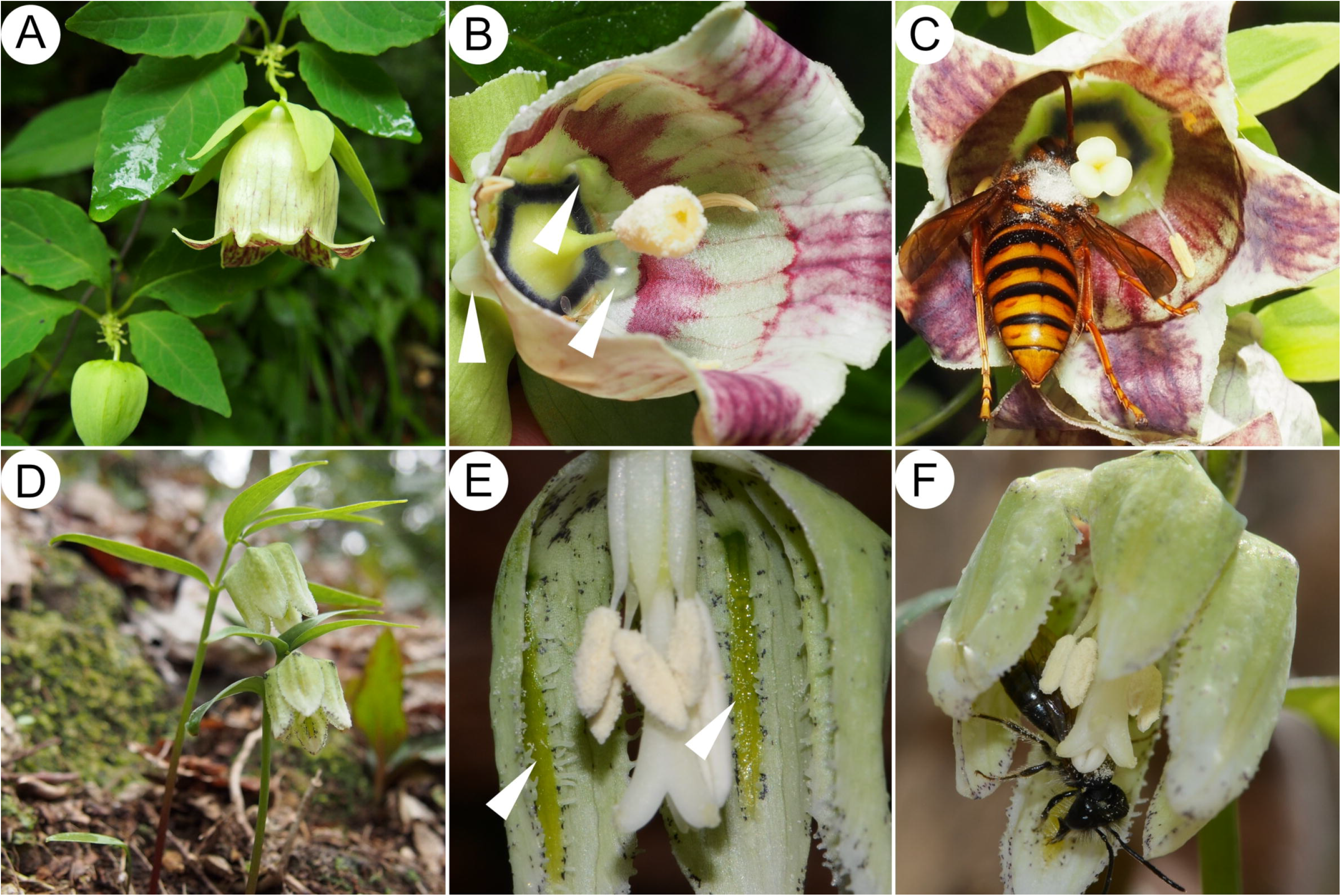
Flowers and pollinators of *Codonopsis lanceolata* (A–C) and *Fritillaria koidzumiana* (D–F). (A) General appearance. (B) Longitudinal section of a flower showing nectar droplets in nipple-shaped spurs (white arrowheads). (C) Pollinator: *Vespa simillima xanthoptera* collecting floral nectar at the base of the flower. Note that its dorsal thorax is dusted with pollen and touches the stigma. (D) General appearance. (E) Longitudinal section of the flower showing band-shaped nectaries on the tepals (white arrowheads). (F) Pollinator: *Andrena* bee visiting a flower with pollen on dorsal thorax. The pollinator manoeuvres itself into the bell-shaped flower.

**Figure 2.**
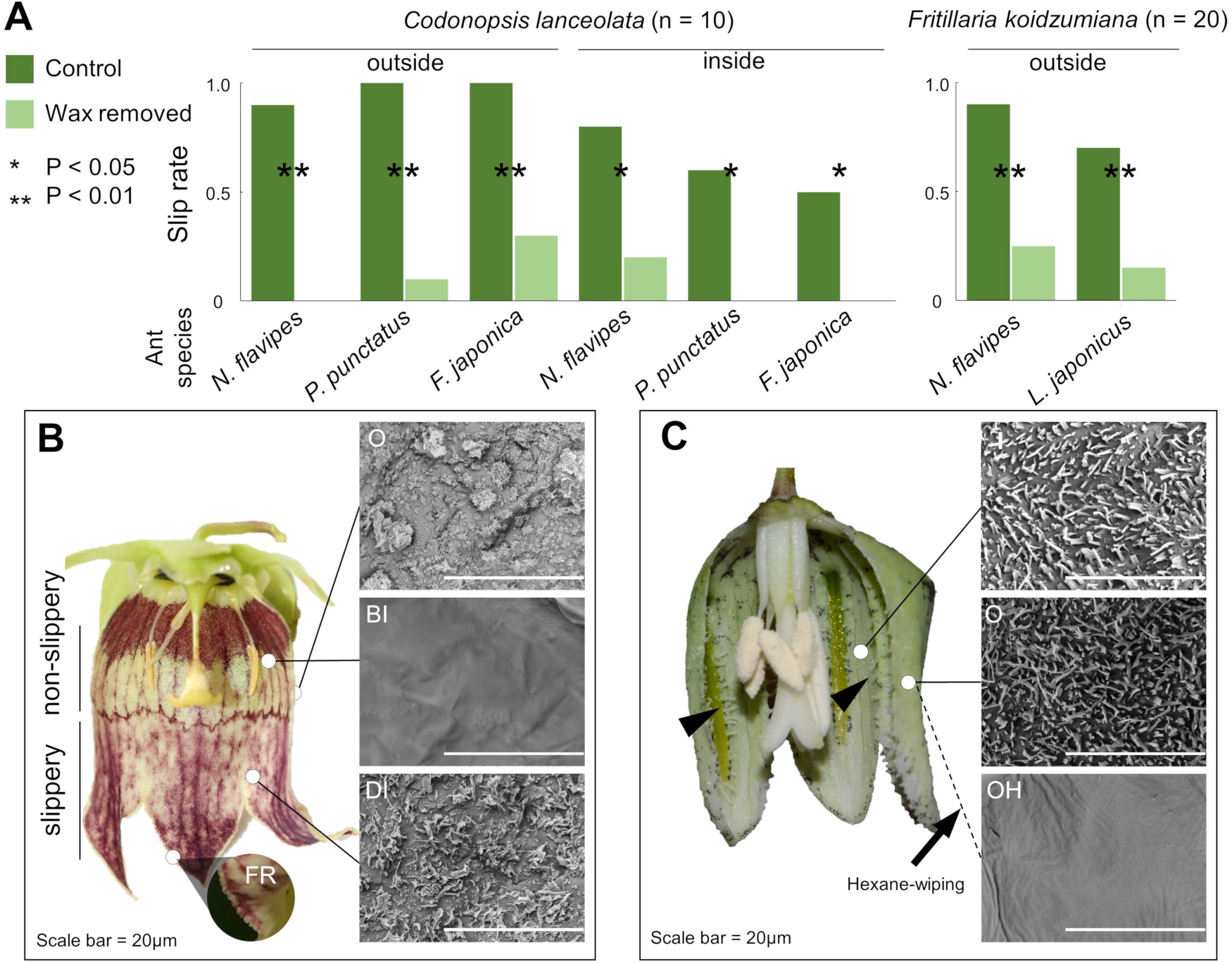
Impact of epicuticular wax crystals on the attachment of freely walking ants. (A) Slip rates of ants in the behavioural assays on *Codonopsis lanceolata* and *Fritillaria koidzumiana* perianths. The dark green bars show the slip rate on the control perianths, and the light green bars show the same on the wax-removed perianths (wax removed by wiping with hexane). The sizes of the ants used in the experiment are: *Nylanderia flavipes*, 2 mm; *Pristomyrmex punctatus*, 3 mm; *Lasius japonicus*, 3 mm; *Formica japonica*, 7 mm. (B, C) Longitudinal sections of the flowers and corresponding scanning electron microscopy (SEM) images of the perianths of *C. lanceolata* (B) and *F. koidzumiana* (C). Arrowheads show the hairy processes on *F. koidzumiana* tepals. O, outer (abaxial) surface of perianths; BI, basal inner (adaxial) surface; DI, distal inner (adaxial) surface; FR, rough structure on the fringe of corolla; I, inner (adaxial) surface: O, outer (abaxial) surface; OH, outer (abaxial) surface wiped with hexane.

## MATERIALS AND METHODS

### Materials

*Codonopsis lanceolata* is a perennial herbaceous vine that occurs along forest edges of evergreen and deciduous forests in temperate areas of East Asia. Each individual produces 1–30 flowers from September to October. The corolla is creamy white with variable degrees of purple patches and lines and a diameter of 23.7 ± 0.7 mm (mean ± SE, n = 6) (Fig. 1A–C). Each flower may hold as much as 94.0 ± 25.1 μL (mean ± SE, n = 3) of nectar during the day (Fig. 1B). As is common to most Campanulaceae, the flower is protandrous. As the bud opens, the pollen on the stamen is deposited and presented on the style column, at which stage the stigma remains unexposed (male stage; Fig. 1B). On the second day, the stigma becomes exposed (female stage; Fig. 1C). The flower withers on the third day. The flowers are almost exclusively visited by hornets (genus *Vespa*; Hymenoptera, Vespidae) that carry abundant pollen on their thoraces (Fig. 1C) and are thus likely effective pollinators (Inoue *et al.* 1990; Funamoto 2019) (K. Takeda, personal observations). We used plant individuals growing at the Ashiu Forest Research Station of Kyoto University (35°18’N, 135°43’E, Kyoto Prefecture, Japan), Sakauchi (35°37’N, 136°22’E, Gifu Prefecture, Japan), and Shiramine (36°10’N, 136°37’E, Ishikawa Prefecture, Japan) for the experiments and observations described below.

*Fritillaria koidzumiana* is a spring ephemeral endemic to central Japan that grows on the forest floors of cool-temperate deciduous forests. Each mature individual produces a single flower at the shoot tip between late March and April. The flower is a broad campanulate, white to cream coloured with green and purple dots, and has a diameter of approximately 12 mm (Fig. 1D-F). Its nectary is band-shaped and located along the centre of each of the six tepals (Fig. 1E). The nectaries are exposed and easily distinguishable by their green colour. The flowers are pollinated by mining bees *Andrena* (Hymenoptera, Andrenidae) (Fig. 1F), mainly *A. benefica* (Naruhashi *et al.* 2006). We used plant individuals from Tsurugi (36°26’N, 136°38’E, Ishikawa Prefecture, Japan) for the analyses presented below.

### Behavioural assay

Previous studies of slippery plant surfaces, such as those in the pitcher plant *Nepenthes* (Nepenthaceae) and the stems of *Macaranga* myrmecophytes (Euphorbiaceae) (e.g. Federle *et al.* 1997; Gaume *et al.* 2002), have shown that the slipperiness is caused by epicuticular wax crystals. Thus, we examined whether epicuticular wax is also responsible for the slipperiness of *C. lanceolata* and *F. koidzumiana* perianths. In the behavioural assays, we used *C. lanceolata* plant individuals from the Ashiu population, and *F. koidzumiana* plant individuals from the Tsurugi population. We first investigated ant behaviour on flowers whose wax had been removed. Because plant epiculaticular waxes are soluble in non-polar solvents, we gently wiped the perianths with glass wool (thickness, 2–6 μm, AS ONE Corporation, Osaka, Japan) soaked with pure hexane (Wako Pure Chemical, Osaka, Japan). We compared the proportions of ants that slipped off the wax-removed flowers to those on control flowers whose perianths were left untreated. Four ant species (Hymenoptera, Formicidae), which have been observed foraging for floral nectar in the natural habitats of the studied plant species, were used for this experiment; *Nylanderia flavipes*, *Pristomyrmex punctatus*, and *Formica japonica* for *C. lanceolata*, and *N. flavipes* and *Lasius japonicus* for *F. koidzumiana*. We carried out experiments on both the adaxial (hereafter, inner or inside) and abaxial (hereafter, outer or outside) surfaces of the corolla for *C. lanceolata,* but only on the outer surface of the tepals for *F. koidzumiana.* This is because *F. koidzumiana* possesses hairy processes on the edges of the outer-whorl tepal and along the boundary of the nectary on the inner tepal surface (Fig. 2C), and ants could walk by gripping them. In the case of *C. lanceolata,* two flowers from each of ten individuals were used for experiments; one was used for the hexane-wipe treatment and the other as a non-treated control. In the case of *F. koidzumiana*, one flower from each of 40 individuals was used for experiment; 20 were treated with a hexane-wipe and 20 were used as non-treated controls. Difference in the sample sizes between the two species simply reflects availability of flowers. We conducted one trial for one individual of each ant species on both sides of every flower. The detailed procedures of the experiment are described in the Supplementary Material.

### Microscopy

We investigated whether epicuticular wax crystals are present on the slippery portions of the perianths by using scanning electron microscopy (SEM). The flowers (*C. lanceolata* from Ashiu, n = 3; *F. koidzumiana* from Tsurugi, n = 4) were sampled from separate individuals in the field, kept in Ziploc® bags (S. C. Johnson & Son, Inc., US) for less than 6 h before being brought back to the laboratory, and frozen at −25°C until examination. Subsequently, the flower samples were freeze-dried overnight using a vacuum freeze dryer (FDU-1200, Tokyo Rikakikai Co. Ltd., Tokyo, Japan), cut into ca. 5 x 5 mm fragments and fixed onto specimen holders with double-sided carbon tape. Although the freeze-drying process makes the perianths slightly shrunken, it does not affect the observation of wax crystals on the perianth surface. The samples were sputtered with gold for 90 sec. (30 mA) using a fine sputter coater (JFC-1200, JEOL Ltd., Tokyo, Japan) and the surface structures were observed using an SEM (TM-3000, Hitachi, Tokyo, Japan). Furthermore, we wiped a portion of the *F. koidzumiana* tepal with hexane-soaked glass wool before freezing, the same treatment as performed in the wax removal behavioural experiment, to compare them to ordinary tepals to determine how the hexane-wiping treatment affected the presence of wax crystals.

### Bridging experiment

To test whether the slipperiness prevents ants from entering flowers under field conditions, we bridged the slippery flower surfaces with non-slippery material and studied the ant behaviour (*C. lanceolata* in the Sakauchi population; *F. koidzumiana* in the Tsurugi population). In the case of *C. lanceolata*, we fixed a piece of masking tape (J7520, Nitoms, Inc., Tokyo, Japan) to span the base of the inner corolla and the floral stalk. As controls, we fixed the tape only to the floral stalk. For *F. koidzumiana*, we used a bamboo stick pinned to the ground beneath the flower, the top of which was attached to the inner surfaces of the tepals (Fig. S1). As controls, we similarly pinned the sticks but not touching the tepals. For *C. lanceolata*, we marked 83 flowers from 18 individuals, and for *F. koidzumiana*, we marked 80 flowers from 80 individuals in the field. We divided the flowers into the treatment group (43 flowers in *C. lanceolata* and 40 flowers in *F. koidzumiana*) and the control group (40 flowers in *C. lanceolata* and 40 flowers in *F. koidzumiana*). We distributed the *C. lanceolata* flowers from each individual evenly into the two groups. We conducted the above treatments in the morning of the first day, and after an acclimatisation time of 2–3 h, we recorded the number and species of ants in each flower every 90 min in *C. lanceolata* and every 60 min in *F. koidzumiana* during the daytime, when the ants were active (09:30 to 16:00). The observations spanned two days in both species. When the flowers withered or bridging materials were accidentally detached from the flowers, we stopped recording and only used the data collected before the accident.

### Ant introduction experiment

Previous studies have reported that nectar-thieving ants attack and deter pollinators, thereby decreasing plant reproductive success (Tsuji *et al.* 2004; Ness 2006; Cembrowski *et al.* 2014). Thus, we artificially introduced ants to *C. lanceolata* flowers in the field to investigate whether ants negatively affect pollination. First, we divided the flowers from each individual evenly into three groups: the ‘ant-present’ treatment group and two control groups. For the ant-present treatment, we secured a live *F. japonica* ant inside the flower by tying the ant with a thread and fixing the other end of the thread to the floral stalk through a hole made on the corolla base (Fig. S1c). For the control flowers, we either only attached the thread (‘thread-only’ control) or left them untreated (‘untreated’ control). In 2017, we only tested the ant-present and thread-only groups, thus the untreated group is only represented in the 2018 data. In total, we used 157 flowers from 25 *C. lanceolata* individuals in this experiment (25 flowers in 2017, 132 flowers in 2018).

To test the effect of ant presence on pollinator hornets, we recorded their behaviours by video camera for 1–4 h during the daytime (9:00–16:00). Specifically, we recorded the species, visit duration, and visitation frequency of hornets. The visitation frequency was calculated based on the number of visits (without discriminating hornet individuals) over 120 min from the start of the observation. Thus, video recordings shorter than 120 min were not used in our visitation frequency calculations.

We then collected all remaining flowers and fruits 19 days after the last flowers had withered and counted the number of fertilised and unfertilised ovules in each fruit, in order to use fruit set and seed set as a proxy for reproductive success. Eight fruits were severely damaged by seed predators and their ovules could not be counted. Hence, we excluded such damaged fruits from the ovule counts. However, if one or more of the three locules remained undamaged, we used data from the undamaged locules.

### Statistical analysis

All statistical analyses were carried out using the R version 3.3.1 (R Core Team 2016). We compared the slip rates in the wax removal experiment between the treatment and control groups using the McNemar test with a binomial distribution in the case of *C. lanceolata* and Fisher’s exact test in the case of *F. koidzumiana*.

To analyse the results of the bridging experiment and the ant-introduction experiment, we assessed the generalised linear model (GLM) using the *glm* function and the generalised linear mixed model (GLMM) using the *glmer* function in the lme4 R package (Bates *et al.* 2015). To compare the frequencies of the ant-present records for each flower treatment, we used GLMM with a binomial distribution and individual ID as a random effect for *C. lanceolata*, and GLM with a binomial distribution for *F. koidzumiana*. We compared the durations between different treatments using GLMM with a gamma distribution, and year, individual ID, and flower ID as random effects. To compare visitation frequencies, we used GLMM with a Poisson distribution, treatments and flower sex stage as explanatory variables, and year and individual ID as random effects. This is because the sex stage of the flower has a large effect on the visitation frequency (male-stage flowers are more attractive), and models including flower sex stage yielded the lowest Akaike information criterion (AIC) values. To compare the fruit set (proportion of flowers that developed as fruits), we used GLMM with a binomial distribution, the fruit set of each flower (1 or 0) as a response variable, and the individual ID as a random effect. To compare the seed set (proportion of ovules that developed as seeds) we used GLMM with a binomial distribution, the seed set of each ovule (1 or 0) as a response variable, and the individual ID and flower ID as random effects.

## RESULTS

### Mechanism of slipperiness

In the behavioural assays, ants were significantly less likely to slip on hexane-wiped flowers than on control flowers in both *C. lanceolata* and *F. koidzumiana* (Fig. 2A, *P* < 0.05). The slippery surfaces of the flowers of the two plant species—the abaxial surface and distal adaxial surface of *C. lanceolata* and both sides of the tepal of the *F. koidzumiana* flowers—were covered by dense epicuticular wax crystals (Fig. 2B, C, S2). In contrast, the basal adaxial surface of the *C. lanceolata* flower, which was not slippery, was not covered by wax crystals. This difference can be seen by the naked eye as a difference in surface optical appearance: the slippery surface has a matt appearance, but the non-slippery surface is distinctly shiny. The boundary between the slippery and non-slippery surfaces is visible as a distinct change in pigment coloration (Fig. 2B, SEM image of the boundary is shown in Figure S2). After wiping with hexane, these crystals were clearly removed from the *F. koidzumiana* tepals (Fig. 2C).

### Ants enter flowers via non-slippery bridges

Nectar-thieving ants included *N. flavipes* (4.0 ± 3.5 individuals per flower, n = 31), *Camponotus umematsui* (1.2 ± 0.44, n = 17) and *F. japonica* (1.0 ± 0.0, n = 5) in *C. lanceolata*, and *F. japonica* (1.0 ± 0.21, n = 23), *L. japonicus* (1.5 ± 0.79, n = 23), and *N. flavipes* (1.6 ± 0.88, n = 9) in *F. koidzumiana* (mean ± SD). In both *C. lanceolata* and *F. koidzumiana*, bridged flowers received ants more often than control flowers (*C. lanceolata*, Fig. 3A, GLMM, Wald test, *P* < 0.001; *F. koidzumiana*, Fig. 3B, GLM, Wald test, *P* << 0.001). Whereas only 10% of *C. lanceolata* flowers and only 5.1% of *F. koidzumiana* flowers in the control group received ants at least once, the percentage increased to 28% in *C. lanceolata* and 45% in *F. koidzumiana* when the flowers were bridged.

**Figure 3.**
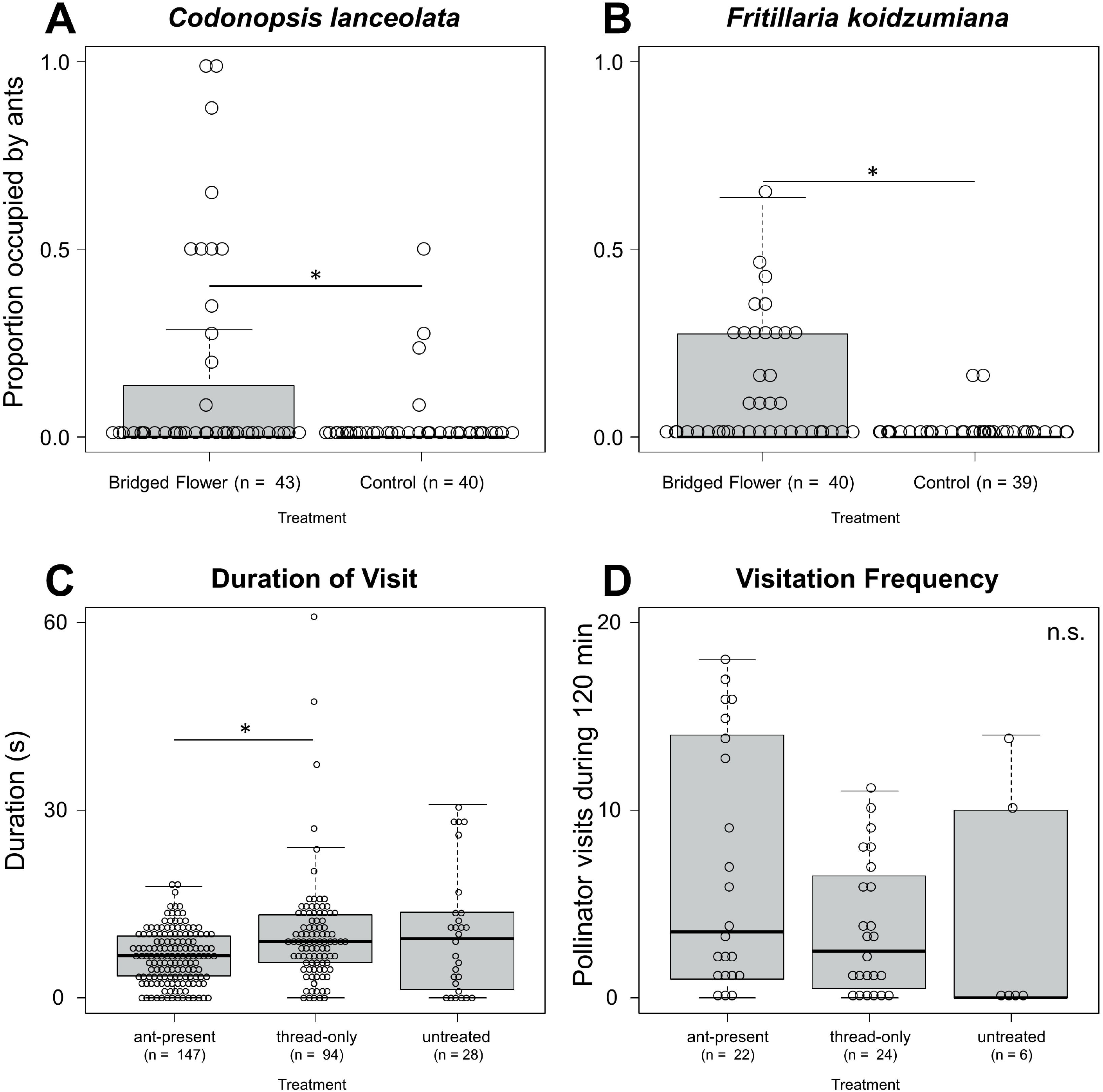
Results of the field experiments. (A–B) Frequencies of the ant-present records for each flower treatment in the bridging experiment. (A) *Codonopsis lanceolata*. (B) *Fritillaria koidzumiana*. The asterisks indicate significant differences according to Wald tests following a generalised linear mixed model (GLMM) or generalised linear model (GLM) (*P* < 0.05). (C-D) Effect of nectar-thieving ants on pollinator behaviour. (C) Duration of pollinator visits to *Codonopsis lanceolata* flowers in the ant-introduction experiment. The asterisks indicate significant differences according to Tukey’s test following generalised linear mixed model (GLMM, *P* < 0.05). (D) Visitation frequency of pollinators to *C. lanceolata* flowers during the 120-min observation period in the ant-introduction experiment.

### Ant effects on pollinators

In total, 306 pollinator visitations were recorded during a cumulative total of 177.2 video recording hours, 268 of which were by *Vespa simillima xanthoptera*, 34 by *V. analis*, and 4 by *Vespa* hornets that could not be identified due to unclear images. After excluding records that were too obscure for us to measure visit durations, we used 269 visitations (147 ant-present, 94 thread-only, and 28 untreated flowers) in the analysis. After alighting on a flower, hornets collected nectar from the five nectar spurs as they moved within the flower, with their dorsal side facing the stylar column (thus contacting the pollen and the stigma; Fig. 1C, Supplementary Movie S3). However, when the hornets came into contact with ants in ant-present flowers, they stepped back abruptly and left the flowers (Supplementary Movie S4). As a result, the hornets stayed at ant-present flowers for significantly shorter durations than at thread-only flowers (Fig. 3C; GLMM, Tukey’s test, *P* = 0.04), but not at untreated flowers (*P* = 0.16). The mean visit duration in ant-present flowers was approximately 65% that of the two control flowers (6.6 s in ant-present, 10.1 s in thread-only, and 10.2 s in untreated flowers). There was no significant difference in visitation frequency among treatments (Fig. 3D; GLMM, Tukey’s test, *P* = 0.73 for control vs. ant-present, *P* = 0.63 for thread-only vs. ant-present, and *P* = 0.97 for thread-only vs. control treatments). Fruit and seed sets were calculated based on 53 fruits resulting from 123 flowers, but no differences were detected among the three treatments (GLMM, Tukey’s test; Table S1; fruit set, *P* = 0.40 for control vs. ant-present, *P* = 0.86 for thread-only vs. ant-present, *P* = 0.72 for thread-only vs. control treatments; seed set, *P* = 0.29 for control vs. ant-present, *P* = 0.77 for thread-only vs. ant-present, *P* = 0.70 for thread-only vs. control treatments).

## DISCUSSION

### Crystals generate perianth slippery properties that deter nectar-thieving ants

The wax removal experiment showed that the hexane-wiping treatment eliminates the slippery properties of the perianths to ants (Fig. 2A). Our SEM observations indicated that the slippery surfaces of *C. lanceolata* and *F. koidzumiana* flowers are densely covered with epicuticular wax crystals, whereas no such crystals were observed on the non-slippery area (basal adaxial surface of *C. lanceolata*) (Fig. 2B, C). On the other hand, there is no difference between the cell shapes of slippery and non-slippery areas; they are both convex or flat in *C. lanceolata* and flat in *F. koidzumiana* (Figure S2). Furthermore, the crystals were removed by hexane-wiping treatment (Fig. 2C). Although we cannot fully rule out the possibility that the hexane-wiping treatment destroyed the epidermal cell surface structure, rather than epicuticular wax per se, a clear association between the presence/absence of wax crystals on slippery and non-slippery portions of *C. lanceolata* corolla indicates that epicuticular wax crystals are most likely responsible for the slippery properties of *C. lanceolata* and *F. koidzumiana* floral surfaces to ants. A number of mutually non-exclusive mechanisms by which wax crystals decrease insect attachment have been proposed: the roughness of the surface decreases the real contact area between insect legs and plant surfaces; easily detachable crystals contaminate the adhesive pads of insects; wax crystals adsorb the pad secretion that reinforces attachment; wax crystals dissolve in pad secretion and thereby thicken the fluid layer (Gorb and Gorb 2017). We suppose a combination of all the hypothesized effects, however, we could not determine which of the above mechanisms is responsible in the two studied plant species.

The bridging experiment suggested that the slipperiness prevents ants from entering flowers, explaining why ants are rarely found in the flowers of these species under natural conditions. When ants are observed on flowers in the wild, such flowers are usually attached to surrounding foliage, which acts as a bridge allowing ants to bypass the slippery area (K. Takeda, personal observation). Although wax crystals may also have other functions, such as controlling transpiration, gas-exchange, or surface temperature (Barthlott 1990; Gorb and Gorb 2017), slippery perianths function as an effective means of deterring ants from flowers in these plants.

Importantly, the slipperiness of the flowers does not necessarily prevent entry by legitimate pollinators (hornets in *C. lanceolata* and andrenid bees in *F. koidzumiana*). This may be due to the presence of ‘footholds’ in these flowers. In *C. lanceolata*, there are non-slippery areas toward the base of the inner corolla (Fig. 2B). In *F. koidzumiana*, there are hair-like processes on the edges of the nectaries and on the edges of the outer tepals (Fig. 2C). When hornets visit *C. lanceolata* flowers, they approach flowers flying, grab the fringes of the petals, which have a rough texture (Fig. 2B), and then reach the non-slippery area with their forelegs (Supplementary Movie S3). Pollinator hornets are large enough (*V. analis*, 24.2 ± 1.7 mm, n = 3; *V. simillima xanthoptera*, 19.8 ± 0.60 mm, n = 5; means ± SE) to stride over the slippery area of the distal adaxial surface (ca. 10 mm), whereas ants cannot, even when they reach the fringe, owing to their small body size (less than 6 mm). Thus, slippery areas and footholds may act in concert to effectively filter out ants while accepting legitimate pollinators.

### Presence of ants in flowers affects pollinator behaviour

In *C. lanceolata*, hornets were disturbed by the presence of ants inside the flowers and consequently remained at flowers containing ants for shorter durations (Fig. 3C). These results are consistent with those of previous studies showing that nectar-thieving ants deter pollinators (Tsuji *et al.* 2004; Ness 2006; Cembrowski *et al.* 2014). Visitation frequency of pollinator hornets did not differ between treatments (Fig. 3D), suggesting that the presence of ants does not have long-lasting effects on the foraging pattern of the pollinators.

In present study, we could not confirm the negative effect of ants on plant reproductive success (seed set and fruit set; Table S1). However, we consider that this result does not necessarily mean that ants in flowers do not have any negative effect. Due to technical limitations, we used only one ant in each flower in this experiment, but ant harassment would likely have a larger effect on pollinator behaviour when flowers are occupied by more than one ant, as is the general pattern in bridging experiments. Introduction of several ants in each flower may be ideal to replicate ant-occupied flowers in natural condition. In addition to the direct effect of ants on pollinator behaviour, depletion of nectar in flowers with ants (Fritz and Morse 1981) may further reduce the visit duration or visitation frequency of pollinators and thus affect pollination success, although we did not evaluate this effect in the present study. Ants may also negatively affect pollination by decreasing the performance of pollen due to antibiotic substances secreted by ant bodies (Galen and Butchart 2003). Further field experiments are needed in order to confirm whether the slippery perianths of *C. lanceolata* and *F. koidzumiana* are adaptive by preventing floral entry by ants.

### Function of slippery petals

A number of studies have proposed that slippery plant surfaces by means of wax crystals act as an ant deterrent mechanism. For example, von Marilaun (1878) reported that the stems of *Salix* trees are covered with wax, which may make the stems slippery and deter nectar-thieving ants. Harley (1991) reported that plants in two Lamiaceae genera, *Hypenia* and *Eriope*, have waxy stems that ants are unable to walk on, and proposed the term ‘greasy pole syndrome’ for the combination of characteristics that seems to prevent ants from climbing stems (Harley 1991). Several other studies have also reported that wax crystals on the stems of various plants can prevent ants from climbing them (Eigenbrode 2004; Whitney *et al.* 2009b; Gorb and Gorb 2011, 2017).

While the examples mentioned above are restricted to wax on stems (Kerner von Marilaun 1878; Harley 1991; Whitney *et al.* 2009b; Gorb and Gorb 2011, 2019) or bracts (Kerner von Marilaun 1878), waxy perianths have rarely been reported (Barthlott 1990) apart from trap flowers (e.g., *Aristolochia* (Aristolochiaceae), *Coryanthes* (Orchidaceae) and *Cypripedium* (Orchidaceae)(Gerlach and Schill 1989; Bänziger *et al.* 2005; Antonelli *et al.* 2009; Oelschlägel *et al.* 2009)). However, Bräuer *et al.* (2017) reported that the perianths of *Lapageria rosea* (Philesiaceae) and *Platycodon grandifloras* (Campanulaceae) have dense wax crystals, and are slippery to insects (honeybees and greenbottle flies). They mentioned that such waxy perianths, like in *C. lanceolata* and *F. koidzumiana*, may deter nectar-thieving ants or other floral antagonists (Bräuer *et al.* 2017).

Apart from wax, floral slipperiness caused by flat perianth epidermal cells has also been proposed to deter nectar-robbing bees (Ojeda *et al.* 2012, 2016; Papiorek *et al.* 2014; Moyroud and Glover 2016). Ojeda *et al.* (2016) examined perianth epidermal structures of related plant species pairs and found that bird-pollinated flowers were more likely to have flat cells on their petal surfaces compared to bee-pollinated relatives, which often have conical cells. Conical cells are proposed to increase the roughness of petal surface and make it easier for pollinators to grip the floral surface (Whitney *et al.* 2009a; Alcorn *et al.* 2012). Thus, the flat cells in bird-pollinated flowers may hinder landing by nectar-robbing bees (Ojeda *et al.* 2016).

Despite the history of studies proposing that slippery plant surfaces may deter unwanted floral visitors, this function has not been explicitly tested by field experiments. Previous studies were based either on behavioural experiment in artificial condition (Whitney *et al.* 2009a; Alcorn *et al.* 2012; Bräuer *et al.* 2017; Gorb and Gorb 2019), simple observation (Kerner von Marilaun 1878; Harley 1991) or comparative analysis (Ojeda *et al.* 2012, 2016; Papiorek *et al.* 2014; Costa *et al.* 2017; Coiro and Barone Lumaga 2018; Kraaij and Kooi 2019). To our knowledge, this is the first study to experimentally demonstrate that slippery plant surfaces prevent floral access by unwanted visitors (ants) in the field. Recent studies on floral defence showed that there is a trade-off between defence and attraction of pollinators (e.g., Barlow *et al.* 2017). If pollinators can discriminate and remember the slipperiness of petal surface (Kevan and Lane 1985; Whitney *et al.* 2009a; Alcorn *et al.* 2012), slippery flowers receive less antagonistic visitors, but may simultaneously attract less pollinators. To elucidate the effect of slippery defence on floral choice by pollinators and the ecological conditions leading to the evolution of slippery defence, further experimental studies, especially those conducted in the field, are needed.

## SUPPLEMENTARY DATA

Supplementary data consist of the following. Details of the wax removal experiment. TableS1: Reproductive success of flowers in each treatment in the ant-introduction experiment. Figure S1: Experimental set-up of the bridging experiment and the ant-introduction experiment. Figure S2: Additional SEM image of petal surface. MovieS1: *Camponotus japonicus* trying to walk on the abaxial surface of a *Codonopsis lanceolata* flower. MovieS2: *Nylanderia flavipes, Aphaenogaster famelica* and *Formica japonica* trying to walk on the adaxial surface of a *C. lanceolata* flower. MovieS3: Visitation behaviour of the pollinator hornet to a *C. lanceolata* flower. MovieS4: Disturbed visitation in an ant-present flower in *C. lanceolata*.

## ACKNOWLEDGEMENTS

We thank Keisuke Koba and Minoru Tamura for supporting our SEM analysis, and Ryosuke Ijichi for supporting us in the field observation.

