## Supplemental Text for "Slippery flowers as a mechanism of defence against nectar-thieving ants"

**Supplemental data**

*Details of behavioural assays*

First, the *C. lanceolata* flowers used in the experiments were detached from plants. Flowers were kept in *Ziploc* bags (S. C. Johnson & Sons, Inc., US) until the assays. The assays were conducted less than four hours after the sampling, when flowers were kept fresh. To test the slipperiness of the adaxial surface of the corolla, we placed single ants at the bottom of upward-facing flowers positioned on the ground and recorded whether they slipped off while climbing up the flowers. To test the abaxial surface of the flowers, we placed ants at the summit of downward-facing flowers and recorded whether they slipped off while walking down the flowers. Ants were recorded as ‘slipped’ either when they fell off or could not walk down the flower in 120 s. This is because ants often stayed still without attempting to walk further on the slippery zone, most of which frequently groomed their feet. This behaviour was not observed on non-slippery surfaces. In the case of *F. koidzumiana*, the flowers were not detached from plants. We placed ants at the summit of downward-facing flowers and recorded whether they slipped off while walking down the flowers in 120 s.

Table S1 Reproductive success of flowers in each treatment in the ant-present experiment.

| Treatment | Fruit set | Seed set (mean ± SD) |
| --- | --- | --- |
| Untreated | 0.46 (n=52) | 0.59 ± 0.22 (n=18) |
| Thread-only | 0.43 (n=37) | 0.51 ± 0.29 (n=14) |
| Ant-present | 0.38 (n=34) | 0.47 ± 0.27 (n=13) |

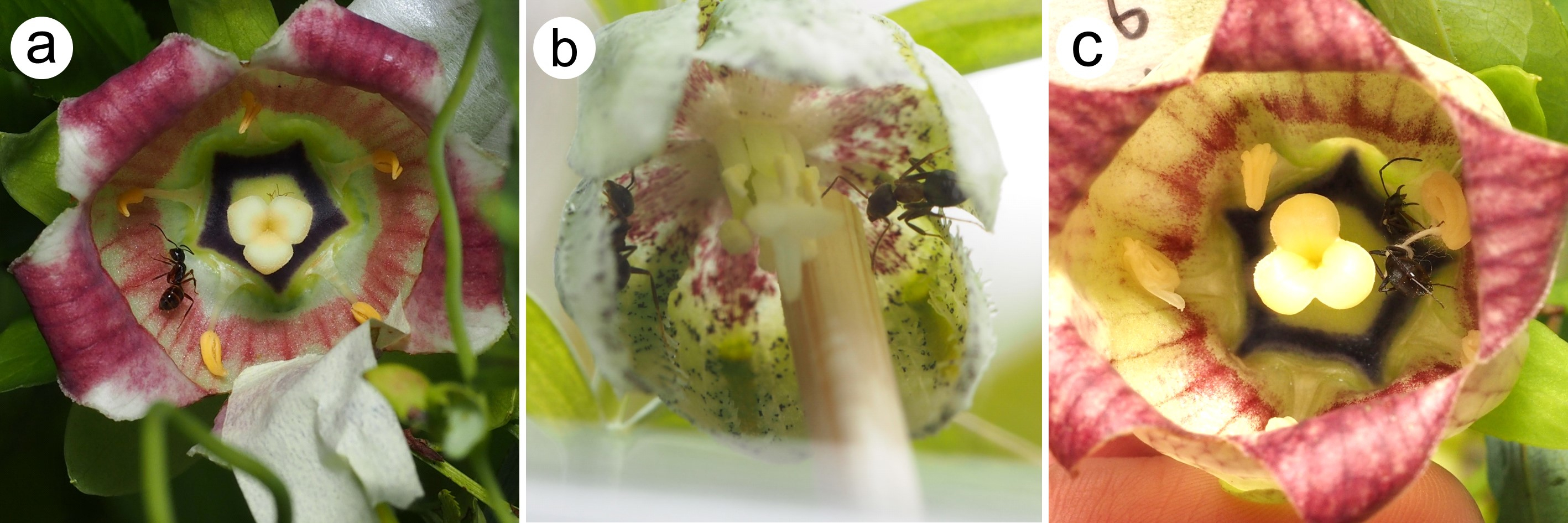

Figure S1

Experimental set-up of the bridging experiment (a, b) and the ant-present experiment (c). (a) The slippery zone of the *Codonopsis lanceolata* flower bridged with masking tape. (b) The flower of *Fritillaria koidzumiana* bridged with a bamboo stick. (c) The ant *Formica japonica* fixed with thread inside the corolla of *C. lanceolata.*

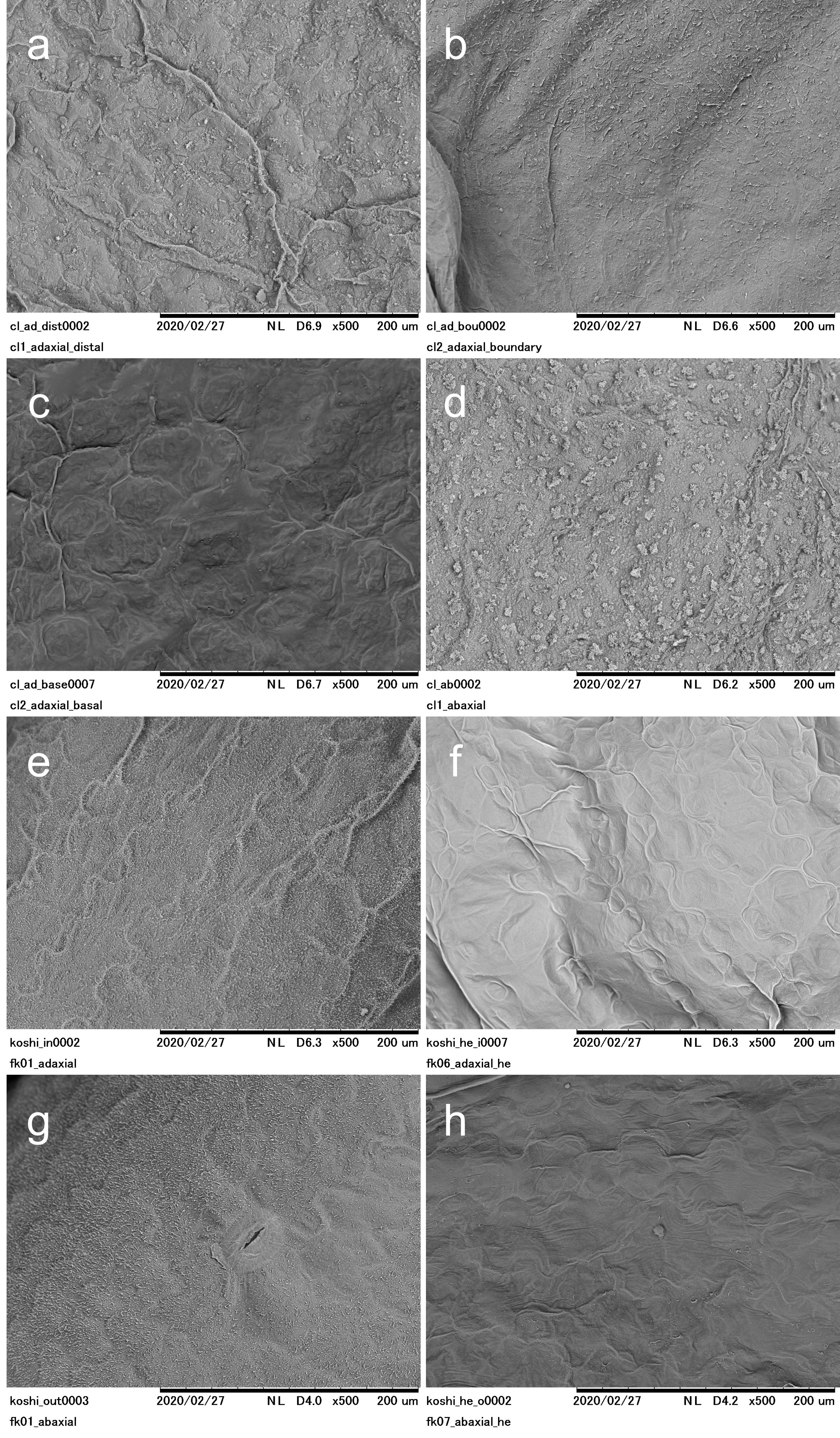
Figure S2

Additional scanning electron microscopy (SEM) images of the perianths of *Codonopsis lanceolata* (a–d) and *Fritillaria koidzumiana* (e–h) under lower magnification.

(a) Distal adaxial surface of *C. lanceolata* petal; (b) boundary of slippery and non-slippery surfaces of *C. lanceolata* petal. The slippery surface is on the upper half, and the non-slippery surface is on the lower half; (c) basal adaxial surface of *C. lanceolata* petal; (d) abaxial surface of *C. lanceolata* petal; (e) adaxial surface of *F. koidzumiana* tepal; (f) adaxial surface of *F. koidzumiana* tepal after hexane-wiping; (g) abaxial surface of *F. koidzumiana* tepal; (h) abaxial surface of *F. koidzumiana* tepal after hexane wiping.
Supplemental movies

S1: *Camponotus japonicus* trying to walk on the outer surface of a *Codonopsis lanceolata* flower. S2: *Nylanderia flavipes*, *Aphaenogaster famelica* and *Formica japonica* trying to walk on the inner surface of a *C. lanceolata* flower. S3: Visitation behaviour of the pollinator hornet to a *C. lanceolata* flower S4: Disturbed visitation in an ant-present flower in *C. lanceolata*.
